# Recombinant Ranpirnase enhances the expression of co-transfected mRNA by modulating cellular metabolic activity

**DOI:** 10.1101/2023.07.22.550185

**Authors:** Xiujin Wu, Liang Chen, Yingxue Lei, Xu Peng, Yaoyi Zhang, Guang Yang, Hailin Yin, Kai Xu

## Abstract

As the world’s first approved prophylactic mRNA vaccine, the development of the Covid-19 mRNA vaccine marks a milestone in mRNA therapeutics. However, achieving efficient transgene expression in therapeutic applications such as protein replacement and cancer therapeutical vaccines remains challenging. This study explores the use of recombinant Ranpirnase, a small ribonuclease, as a novel expression-enhancing tool for mRNA-based therapeutics. In Balb/c mice, co-expression of Ranpirnase significantly increased transgene expression levels by 3-to 6-fold and prolonged expression duration by more than 85%, regardless of mRNA nucleoside modification status. Recombinant Ranpirnase induces a cytostatic state in host cells, stabilizing mRNA and protein transcripts, thereby enhancing transgene expression. Importantly, co-expression of Ranpirnase recombinants did not cause detectable cytotoxicity or alter the tissue distribution of transgene expression, ensuring safety and tolerability at higher translation levels. The compact size of the recombinant Ranpirnase gene allows for seamless fusion with the target gene or co-transfection with existing delivery technologies. Altogether, recombinant Ranpirnase shows promise as an expression-enhancing element in mRNA-based therapeutics, including veterinary applications.

## 1. Introduction

Covid-19 vaccines, Comirnaty (BioNTech/Pfizer) and Spikevax (Moderna), represent the world’s first officially approved prophylactic mRNA vaccines^1–4^, signifying a significant advancement in mRNA therapeutics^5^. While prophylactic vaccines have paved the way, therapeutic applications such as protein replacement therapy and cancer therapeutic vaccines often require higher expression levels and longer durations of transgene expression. However, achieving effective transgene expression is hindered by robust cellular defense mechanisms, including RNA decay mechanism and the innate immune response of host cells. Overcoming these challenges remains a primary focus in mRNA research^6^.

The mRNA molecule, synthesized through an in-vitro transcription (IVT) reaction, triggers the innate immune system through Toll-like receptors (TLRs) and retinoic acid-inducible gene-I (RIG-I)-like receptors^7, 8^. Furthermore, double-stranded RNA (dsRNA) and uncapped 5-triphosphate motifs can also stimulate an innate immune response^9^. A significant breakthrough in mRNA technology has been the incorporation of modified nucleosides during IVT production, effectively suppressing mRNA immunogenicity^10^. Modified nucleosides like N1-methylpseudouridine (mѱ), pseudouridine (ѱ), and N-6-methyladenosine (m6A) have been employed to bypass innate immune activation in host cells^7, 8, 11, 12^. Additionally, the elimination of dsRNA and 5-triphosphate motif impurities further reduces the innate immune response^9, 13^. The mѱ-modification enhances mRNA stability and translation efficiency, providing a solid foundation for the success of the COVID-19 mRNA vaccine. Comparative studies have demonstrated the superiority of mѱ-modified mRNA vaccines over other vaccine types^10^.

Another strategy involves inhibiting the biological activity of host cells, thereby suppressing their innate immune reactivity and RNA decay capacity. Co-administration of low-molecular-weight immunosuppressive molecules, such as an integrated stress response inhibitor and dexamethasone, during mRNA-LNP transfection has shown promising results in increasing protein expression in mice^14^. Similarly, the co-delivery of DNA/siRNA nanoparticles targeting transcription factors involved in inflammatory response pathways has prolonged the duration of transgene expression^15^. However, implementing these strategies remains challenging.

Ranpirnase, a small ribonuclease enzyme, was discovered in the oocytes of the Northern Leopard Frog (Rana pipiens)^16, 17^. It shares 30% amino acid sequence identity but has a similar structural fold to RNase A^17, 18^. Ranpirnase demonstrates unique ribonucleolytic activities by selectively targeting tRNA^19^ and miRNA precursors^20, 21^ while evading the ubiquitous ribonuclease inhibitor (RI)^22, 23^ in eukaryotic cells, resulting in reduced protein translation efficiency, metabolic activity, and innate immune response. Ranpirnase, aka Onconase, has demonstrated cytotoxic and cytostatic effects on tumor cells and has been investigated as a potential treatment for cancer and viral infections^24, 25^.

The antitumor cytotoxicity of Ranpirnase is attributed to its N-terminal modifications, namely glutamate pyrophosphorylation and cyclization between the first and ninth amino residues^18, 26^. However, recombinant Ranpirnase lacking these modifications exhibits reduced cytotoxicity and antigenicity while retaining its ribonucleolytic activity^19, 27^ and RI evasion capability^28^. The biological function of amphibian ribonucleases contributes to reducing the metabolic activity and innate immune reactivity of host cells, inducing a cytostatic state^16, 19, 29^. This cytostatic state may stabilize mRNA and its protein transcripts, resulting in increased transgene expression levels and duration.

This study aimed to investigate the effect of recombinant Ranpirnase co-expression on transgene expression. The results indicated that the presence of recombinant Ranpirnase significantly increased transgene expression in Balb/c mice, resulting in a three-to six-fold increase and prolonging the expression duration by over 85%, regardless of the nucleoside modification status of the mRNA. In vitro studies suggests that the cytostatic effect of recombinant Ranpirnase as a potential mechanism for enhancing gene expression. Notably, these effects were achieved through reduced cellular metabolism without altering tissue distribution of transgene expression or causing any detectable cytotoxicity.

## 2. Material and Method

### 2.1. Animal

The animal experiments were conducted at the Laboratory Animal Center of Sichuan University, in accordance with the National Institutes of Health Guide for Animal Welfare in China. Approval for the study was obtained from the Institutional Animal Care and Use Committee of Sichuan University. Female Balb/c mice, aged 6-8 weeks and weighing between 18-20 grams at the start of the study, were bred in-house. Prior to the experiment, the mice were acclimated for one week and then randomly assigned to treatment and control groups.

For blood collection and serum preparation, mice were anesthetized and blood was collected from the orbital sinus using a vacutainer with heparin anticoagulant. The collected blood was stored at 4°C for two hours, followed by centrifugation at 1,000 x g for ten minutes. The resulting supernatants were transferred into new tubes and stored for long-term preservation at -70°C or for up to two months at -20°C.

### 2.2. Serum biochemical and blood analyses

Serum biochemistry analysis was conducted at Huaxi Ruipai Pet Hospital using a veterinary clinical biochemical detector manufactured by IDXX. Blood routine testing was performed using VetScan HM5 (Abaxis).

### 2.3. mRNA synthesis and plasmid preparation

The CovS mRNA is identical to the BNT162b2 mRNA (Pfizer/BioNTech) and contains specific codon region substitutions for a prefusion version of the SARS-CoV-2 Spike protein with G614D, K986P, and V987P substitutions. The mRNA synthesis was carried out using an in-vitro transcription reaction as previously reported^30^.

### 2.4. LNP and LPX Preparation and Characterization

LNPs were prepared and characterized as previously reported. LNP^⊕^74, a modified LNP with additional Dotap, was used to eliminate systemic delivery^30^. An 8 mM Dotap:Lecithin liposome (DL) was prepared from Dotap/Lecithin mixture at a molar ratio of 20:9 by a thin-film evaporation protocol reported previously^31^. Empty DL (eDL) was obtained by mixing an equal volume of DL with D5W. DNA:DL complexes were prepared by mixing equal volume of DNA (0.85 µg/µl) and DL (8 mM relative to Dotap).

### 2.5. Bioluminescent imaging

Bioluminescent imaging (BLI) was performed using a Xenogen IVIS-200 imaging system, as reported in the reference. The total amount of accumulated Fluc protein over time post-injection was calculated using the AUC analysis, as described by Pardi et al^32^.

### 2.6. Immunization

An immunization experiment was conducted on Balb/c mice, consisting of primary and booster immunizations spaced three weeks apart as reported^30^. Plasma samples were collected at 1, 2, and 3 weeks after the booster immunization to determine spike-specific IgG titers and competitive RBD-ACE2 binding IgG titers.

### 2.7. Enzyme-Linked Immunosorbent Assay (ELISA)

Plasma samples were analyzed using an indirect ELISA protocol, as previously reported^30^.

### 2.8. The TNF-α and IFN-γ ELISA

The concentrations of serum TNF-α and IFN-γ were determined using Mouse IFN gamma Uncoated ELISA and Mouse TNF alpha Uncoated ELISA (Invitrogen), following the protocols recommended by the vendor.

### 2.9. Surrogate ELISA for RBD-ACE2 competitive neutralizing antibodies

A surrogate ELISA was performed as previously reported^33^. Modification was made with the replacement of ELISA Coating Buffer with 1% non-fat milk in PBS buffer. The titer of competitive RBD-ACE2 binding antibody was calculated according to the reported method.

### 2.10. Cell Viability Assay

The metabolic activity of cells treated with ribonuclease recombinants was assessed using an XTT assay (Roche Molecular Biochemicals) as described previously^34^. NCI-H1299 cells were seeded in 96-well microplates at 5,000 cells per well and transfected with different amounts of DNA:DL complexes. Cell viability was calculated as the percentage of absorbency of treated cells relative to eDL-treated control cells. The experiments were repeated twice with triplicate samples for each treatment in each individual experiment. GFP and RFP expression were examined and photographed. Supplementary File X provides details of a pilot study conducted to identify an appropriate DNA dose with minimal cytotoxic effects. Based on the results of the pilot study, a DNA dose of 42.5 ng per well was determined to be suitable for subsequent metabolic evaluations.

### 2.11. Clonogenic Assay

Colony formation inhibition by Ranpirnase recombinant DNA was examined in A549, NCI-H322, and NCI-H1299 cells. DNA:DL complexes were prepared by mixing the DNA mixture with DL liposome. Cells were seeded in triplicates in 6-well plates and transfected with the DNA:DL complexes in the presence of 10% FBS for 24 hours. The transfected cells were then cultured under G418 selection until colony formation. Cells were fixed in Formalin, stained with Giemsa stain, and photographed. The colony formation inhibition rate was calculated relative to eDL-treated cells.

### 2.12. Histology Analysis

Mice (n = 4) were intramuscularly injected with 50 µL of mRNA-LNP containing 15 µg of mѱ-modified CovS and Fluc mRNA, including the rRanp fused version. After 48 hours, the mice were sacrificed, and liver and muscle tissues at the injection site were collected. Tissues were sectioned, stained with H&E, and blindly graded by the lead investigator under guidance of a board-certified pathologist using a modified Allred scoring system^35^ based on severity for myofiber necrosis/degeneration, mixed-cell infiltration, and interstitial edema in the liver and muscle.

### 2.13. Statistical Analyses

Experimental data were statistically analyzed using IBM SPSS Statistics v20. One-Way ANOVA with Bonferroni post-hoc tests was applied, with a significance level set at *P≤*0.05.

## 3. Result

Motivated by the clinical advancements in Ranpirnase against malignant mesothelioma and antiviral therapy, we conducted in vitro investigations on recombinant Ranpirnase and its analogs. Contrary to initial expectations, recombinant Ranpirnase did not exhibit cytotoxicity; instead, it enhanced the expression of co-transfected reporter genes. To assess the expression enhancement of representative members from the RNase A family, namely Ranpirnase, Amphinase-1^36^, BS-RNase^37^, and RNase I, their recombinants (rRanp, rAmph-1, rBS-RNase, and rRNase I) were evaluated in vivo. A Ranpirnase K31R mutant (rRpK31R) with reduced ribonucleolytic activity^25^ was also included for comparison.

### 3.1. Transgene Expression Enhanced by Recombinants of RNase A family Members

We designed a dual-gene mRNA molecule using a Fluc mRNA sequence^30^, with the recombinant ribonuclease gene fused in-frame at the C-terminal of the Fluc gene, separated by a self-cleaving P2A fragment (Fig 1a). Recombinant sequences are compared in Supplementary Figure 1. Balb/c mice received 5 µg of mѱ-mRNA-LNP intramuscularly, followed by bioluminescence imaging (BLI) analysis (Fig 1b). Co-expression of rRanp, rRpK31R, and rAmph-1 significantly increased Fluc expression in the liver and at the injection site compared to the control group. Noticeable Fluc signals were detected on day 13 post-injection and the enhancement of overall expression was compared by using the Area Under the Curve (AUC) of Fluc flux (Fig 1c). Co-transfection of Fluc with rRNase I showed no statistical difference compared to the control, while rBS-RNase demonstrated moderate expression enhancement. The half-life of Fluc activity correlated well with its gene expression enhancement capacity (Fig 1d).

**Fig.1.**
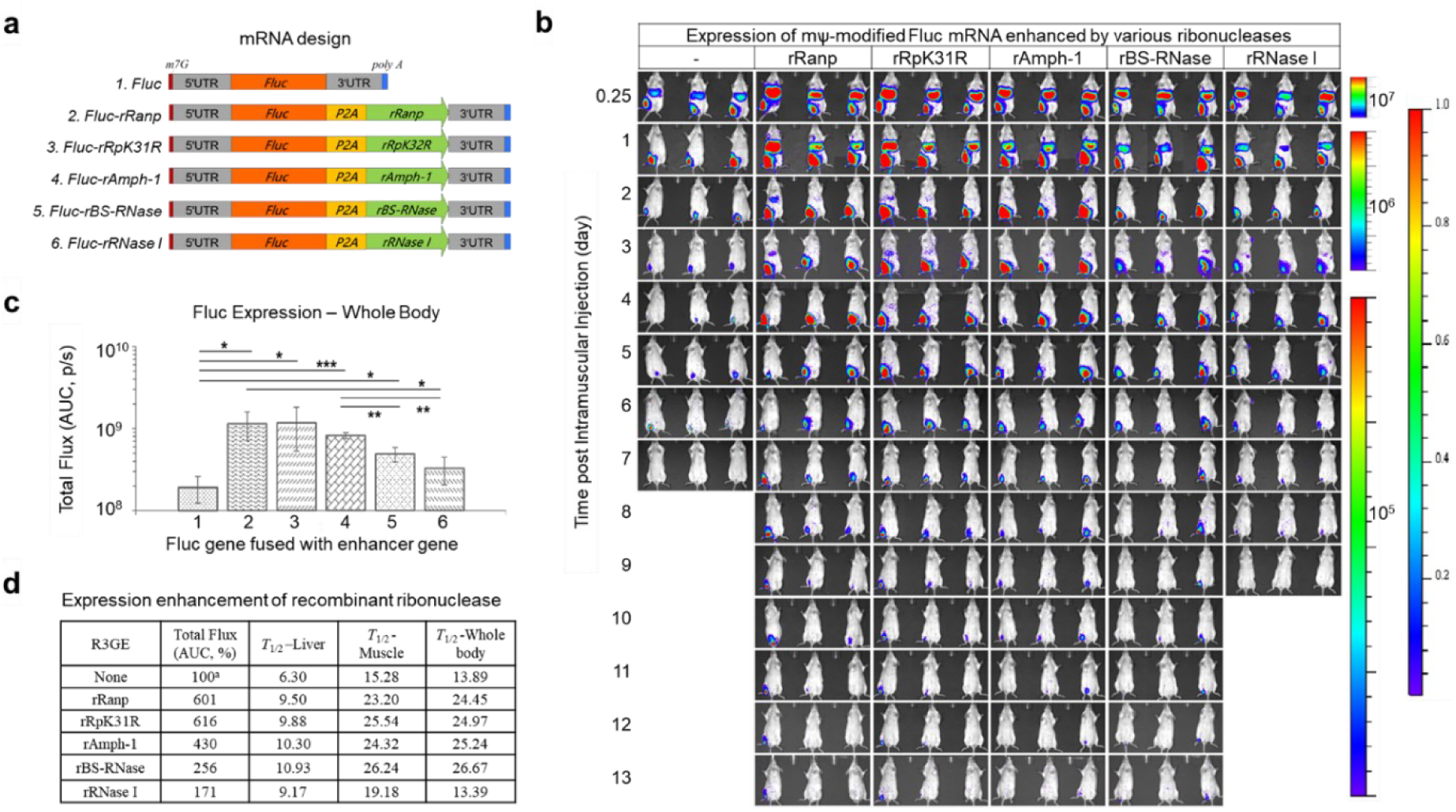
Enhancement of Fluc mRNA Expression by Ribonuclease Recombinants. (a) Schematic representation of the structure and design of Fluc mRNA fused with ribonuclease recombinants. (b) Bioluminescent images of mice obtained using IVIS imaging at various time points (0.25, 1 to 13 days) following intramuscular injection of 5 µg of mѱ-modified mRNA encapsulated in LNP (n = 3 mice per group). (c) Quantification of the expression enhancement by 5 µg of mRNAs: 1. Fluc, 2. Fluc-rRanp, 3. Fluc-rRpK31R, 4. Fluc-rAmph-1, 5. Fluc-rBS-RNase, and 6. Fluc-rRNase I, by using the AUC of Fluc flux of whole-body between 0.25 to 13 days. (d) Half-life of Fluc activity between 0.25 to 13 days in liver, muscle, and whole-body tissues respectively. Flux cutoff: 1x10^4^ p/s/cm^2^/sr. Statistical significance between cohorts was determined using One-Way ANOVA with Bonferroni post-hoc tests (\**p* <0.05, \*\**p* < 0.01 and \*\*\**p* < 0.001). Error bars represent mean ± SEM.

Amphibian ribonucleases exhibit weak but selective ribonucleolytic activity and RI evasion compared to the robust activity of RNase I. These unique features are shared by the ribonuclease recombinants that enhance gene expression. In multiple experiments, co-transfection with rRanp consistently led to a three-to six-fold increase in Fluc activity across tissues, without altering tissue distribution. These findings suggest that the rRanp approach holds considerable potential for universally augmenting gene expression.

### 3.2. The Expression of mѱ-Modified mRNA Enhanced by rRanp Co-Expression

To examine the impact of rRanp on the expression of mѱ-modified mRNA, mѱFluc and mѱFluc-rRanp mRNA were produced and encapsulated in LNP and LNP^⊕^74^30^. Balb/c mice were intramuscularly administered with mRNA-LNPs, followed by BLI analysis (Supplementary Figure 2, and Fig 2ab). The overall Fluc expression in mice treated with rRanp co-expressed mRNA was 3.9-fold higher than that of the mѱFluc-LNP control group (Fig 2c), while the Fluc expression in the liver and muscle of mice was 6.1- and 1.6-fold higher, respectively. Moreover, co-expression of rRanp significantly extended Fluc expression duration by 50.8% and 51.8% in half-life observed in the liver and muscle, respectively (Fig 2ab).

**Fig.2.**
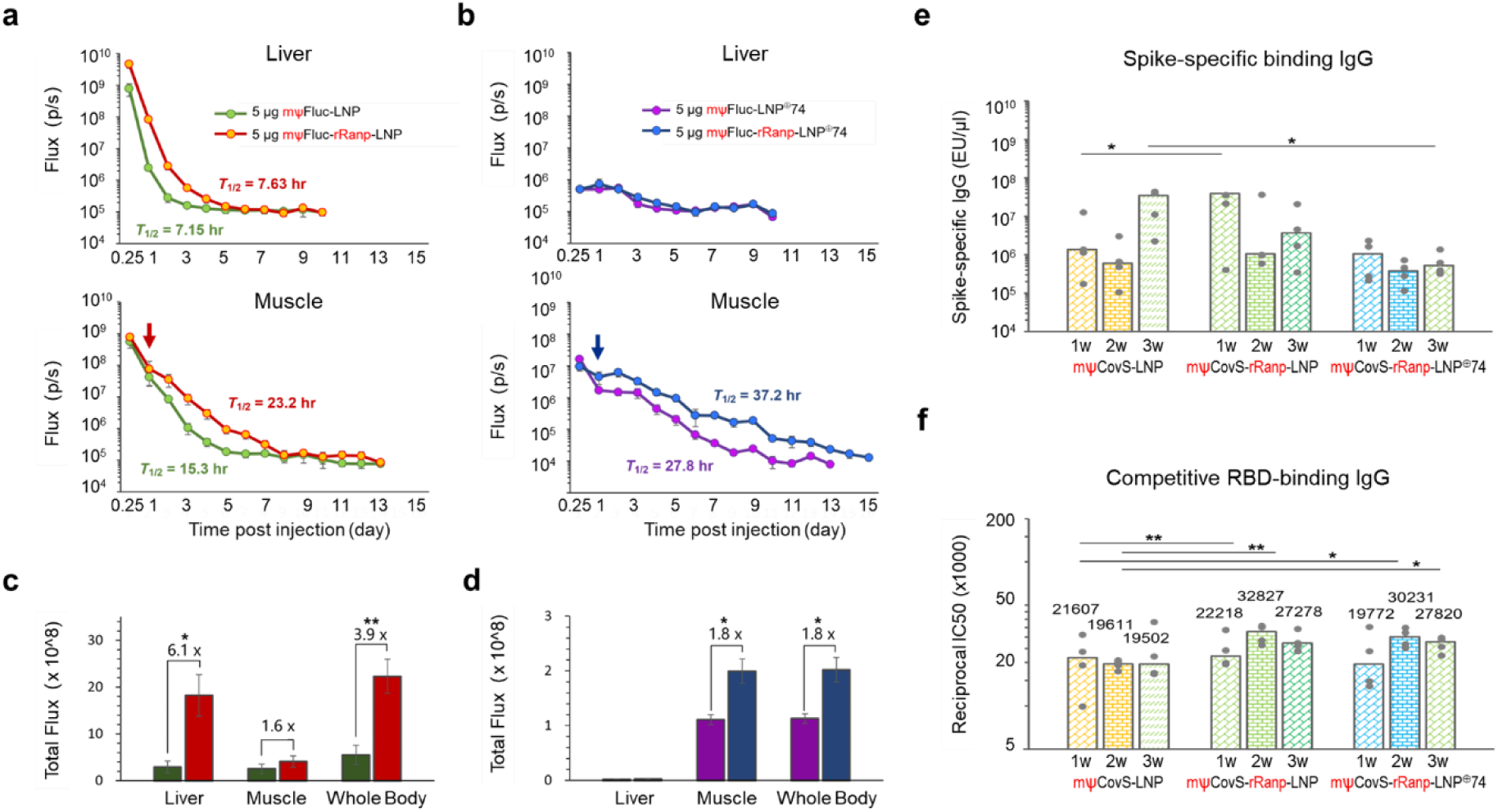
Enhancement of mѱ-Modified mRNA Expression by rRanp Co-Expression. Flux levels in liver and muscle tissues (n = 3 mice per group) measured at multiple time points (0.25, 1 to 15 days) following intramuscular injection of mѱFluc and mѱFluc-rRanp mRNAs encapsulated in LNP (a) and in LNP^⊕^74 (b). (c) Total Fluc flux in liver, muscle, and whole-body tissues of mice that received mѱFluc or mѱFluc-rRanp mRNAs encapsulated in LNP. (d) Total Fluc protein expressed in liver, muscle, and whole-body tissues of mice that received mѱFluc or mѱFluc-rRanp mRNAs encapsulated in LNP^⊕^74 represented by the AUC. Data presented in color code. Flux cutoff: 1x10^4^ p/s/cm^2^/sr. (e) IgG titer determined by Spike-protein specific binding antibody ELISA following immunization of Balb/c mice (n = 4 per group) with 5 µg of mѱCovS mRNA or mѱCovS-rRanp mRNA encapsulated in LPNs. Plasma samples were collected after booster immunization. (f) Reciprocal IC50 titer assessed by competitive RBD-ACE2 binding ELISA. Each dot represents an individual animal. Statistical significance between cohorts was determined using One-Way ANOVA with Bonferroni post-hoc tests (\**p* < 0.05, \*\**p* < 0.01). The negative control group is not presented.

The Fluc expression in the liver was largely eliminated by LNP^⊕^74 formula regardless of rRanp presence, and Fluc expression kinetics in the muscle was significantly improved in the presence of rRanp delivered by LNP^⊕^74 (Supplementary Fig 2, Fig 2bd). At 24 hours after injection, a distinct turning point (arrow) was observed, reflecting the effect of rRanp activity, and higher expression levels were maintained afterward. The overall Fluc expression increased 1.8-fold in muscle, with the half-life of Fluc expression improved from 27.8 to 37.2 hours, a 33.9% increase over the mѱFluc-LNP^⊕^74 control (Fig 2d).

To evaluate the potential impact of rRanp enhancement on the immunogenicity of mѱ-modified mRNA vaccines, an rRanp-enhanced version of CovS mRNA^30^ was constructed based on the framework depicted in Figure 1a. An immunization experiment consisting of primary and booster immunizations spaced three weeks apart was conducted, and immunogenicity was measured using a modified ELISA. After the booster immunization, comparable levels of spike-specific IgG titers were detected in samples from all vaccines (Fig 2e). The competitive RBD-ACE2 binding IgG titers were significantly higher in rRanp-enhanced groups (Fig 2f) regardless the LNP formulation. The LNP^⊕^74 vaccine achieved the same level of immunity by unitizing 9% antigenic proteins produced in the LNP vaccine controls when protein expression was judged by Fluc expression (Fig 2cf). Both vaccines elicited more immunity than the mѱCovS-LNP control vaccine. Regardless of the delivery formular, the protein expression kinetics promoted by rRanp enhancement were more consistent with the exponential-incremental dose principle for effective antibody production^38^. The results indicate that the protein produced in the presence of rRanp was antigenic.

### 3.3. The Expression of unmodified mRNA Enhanced by rRanp Co-Expression

To evaluate the enhancement of gene expression by rRanp on unmodified mRNA, Balb/c mice were intramuscularly injected with modified and unmodified mRNA encapsulated in LNP. The results of the BLI analysis are presented in Figure 3abc. The overall Fluc expression was significantly higher in mice treated with rRanp-enhanced mRNA compared to its unmodified mRNA counterpart, although lower than that produced by mѱ-modified Fluc mRNA (Fig 2ac). Owing to the significant initial decrease in expression of unmodified mRNA, the overall Fluc expression by rRanp-enhanced mRNA at doses of 5 µg and 10 µg was only 50% and 85% of that of mѱ-modified Fluc mRNA, respectively. However, the rRanp-enhanced expression kinetics paralleled or exceeded that of mѱ-modified mRNA at different time points in the liver and muscle. The expression kinetic was improved further when mRNA dosage was increased (Fig 3ab), providing an alternative approach for vaccine development. A notable inflection point in the changes of expression kinetics was observed in muscle 1 day after injection (Fig 3ab, arrows in muscle). By excluding the initial drop in expression right after injection, rRanp improved the expression kinetics of unmodified mRNA over that of mѱ-modified mRNA.

**Fig.3.**
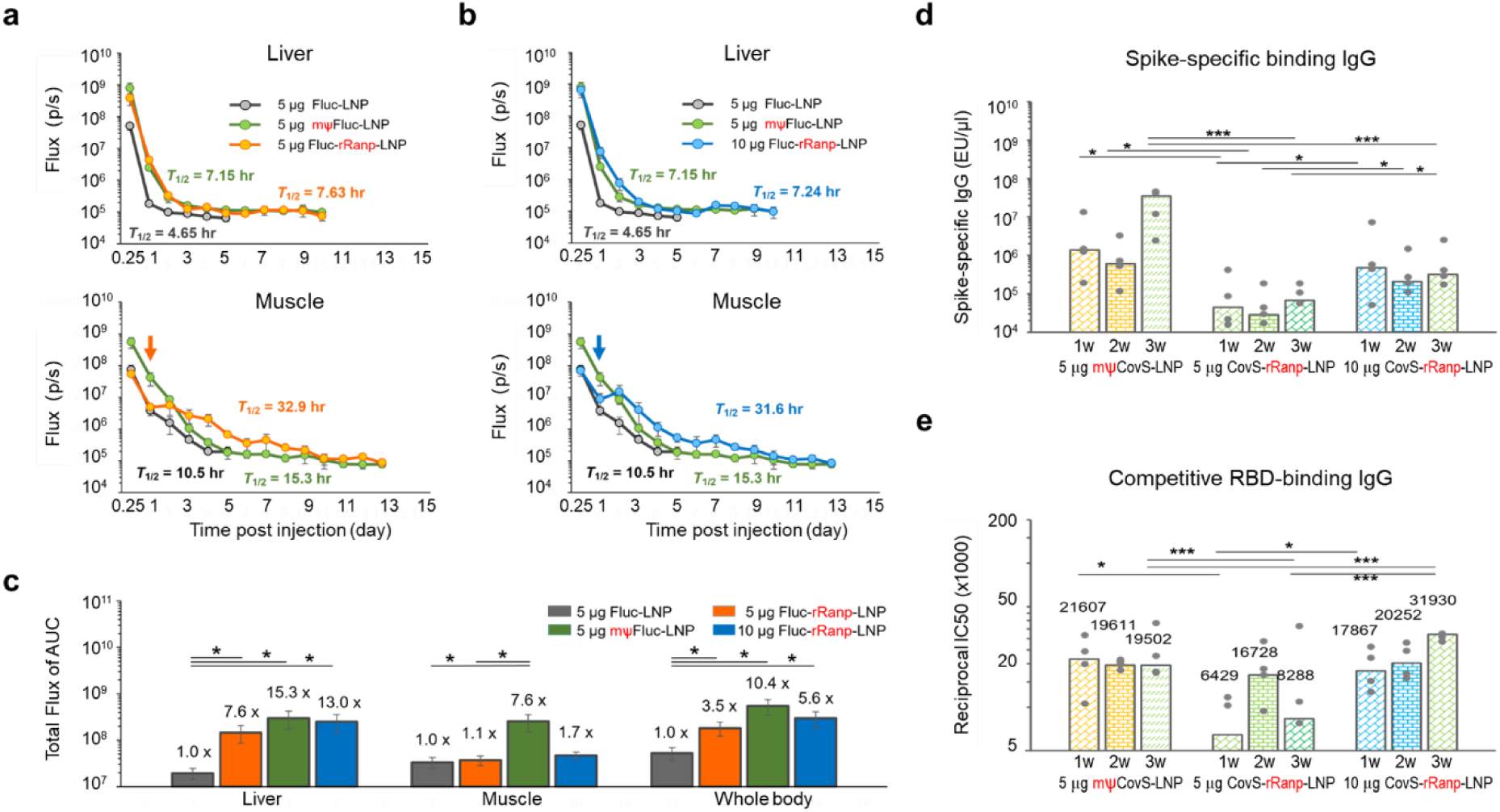
Enhancement of Unmodified mRNA Expression by rRanp Co-Expression. Flux levels in liver and muscle tissues at multiple time points (0.25, 1 to 13 days) after intramuscular injection of 5 µg of Fluc-rRanp mRNA (a) and 10 µg of Fluc-rRanp mRNA (b), respectively, compared to Fluc control mRNA and mѱFluc control mRNA, both encapsulated in LNP. Flux cutoff: 1x10^4^ p/s/cm^2^/sr. (c) Total Fluc protein expression in liver, muscle, and whole-body tissues represented by the AUC. (d) IgG titer determined by Spike-protein specific binding antibody ELISA following immunization of Balb/c mice (n = 4 per group). Plasma samples were collected at weeks after booster immunization. (e) Reciprocal IC50 titer assessed by competitive RBD-ACE2 binding ELISA. Each dot represents an individual animal. The dashed line indicates the assay’s detection limit. Statistical significance between cohorts was determined using One-Way ANOVA with Bonferroni post-hoc tests (\**p* < 0.05, \*\**p* < 0.01, \*\*\**p* < 0.001). Error bars represent mean ± SEM. The negative control group is not presented.

To assess the effect of rRanp on the immunogenicity of unmodified mRNA vaccine, the earlier immunization experiment was replicated with different doses of CovS-rRanp-LNP vaccine. At a dose of 5 µg of mRNA, a marked reduction in spike-specific IgG titer was detected in plasma samples from CovS-rRanp-LNP vaccine group compared to that of mѱCovS-rRanp-LNP control vaccine after booster immunization (Fig 3d). However, the measurements of competitive RBD-ACE2 binding IgG titers from mice treated with 10 µg of CovS-rRanp-LNP exhibited significantly improved immunogenic kinetics over that of the mѱ-modified CovS vaccine (Fig 3e).

### 3.4. rRanp enhanced CMV-driven plasmid expression through cytostasis in vitro

The CMV-driven rRanp expression plasmid, based on the pEGFP-c3 framework, was constructed and evaluated for its effect on the expression of co-transfected plasmid DNA in cultured cells. pEGFP-c3 plasmid DNA was mixed with Fluc-, rRanp-, and p53-expression plasmid DNA at a ratio of 1:9 and encapsulated in DL liposomal complexes. DNA-DL complexes were transferred into NCI-H1299, NCI-H322, and A549 cells, and green fluorescence was observed for 6 days. Representative images of NCI-H1299 and the others are displayed in Fig 4a and Supplementary Figure 3. The Co-transfection of rRanp significantly increased overall GFP levels and expression duration in all tested cells (Fig. 4b), while p53 co-expression inhibited eGFP expression.

**Fig.4.**
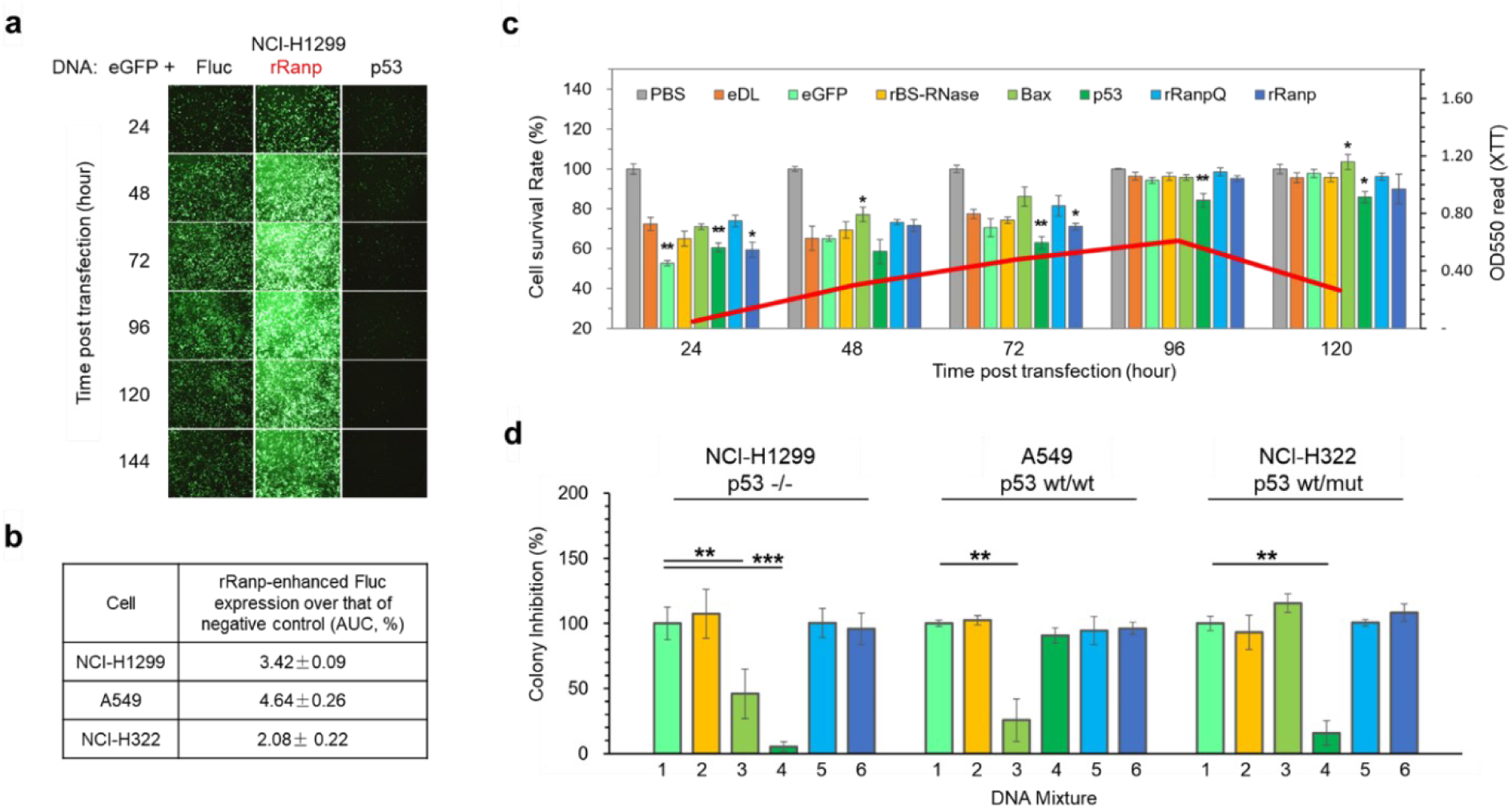
Effect of Ribonuclease Recombinants on Gene Expression and Toxicity in vitro. (a) Influence of co-transfection of Fluc, rRanp, and p53 DNAs on the expression of pEGFP-c3 in NCI-H1299 cells. (b) Enhancement of Fluc protein expression quantified by AUC of flux in cultured cell lines. (c) Altered metabolic activity resulting from transfection with DNA:DL complexes containing: eDL, eGFP, BS-RNase, Bax, p53, rRanpQ, and rRanp plasmids, measured using an XTT assay. (d) Suppression of colony formation by co-transfection of pcDNA3.1 DNA mixed with the following plasmid DNA constructs: 1. eGFP, 2. BS-RNase, 3. Bax, 4. p53, 5. rRanpQ, and 6. rRanp in cultured cells. Statistical significance between cohorts was assessed using One-Way ANOVA with Bonferroni post-hoc tests (\**p* < 0.05, \*\**p* < 0.01, and \*\*\**p* < 0.001). The error bars represent the mean ± SEM.

Metabolic activity of NCI-H1299 cells treated with recombinant ribonucleases was assessed using an XTT assay^34^. DNA mixtures of pEGFP, pRFP, and Ranpirnase recombinant plasmids (1:1:8 ratio) were prepared. DNA:DL complexes were formed by mixing DNA with DL. A pilot study determined a non-toxic pEGFP:DL complex dose of 42.5 ng (Supplementary File). Cells treated with PBS showed exponential growth and served as the reference (Fig 4c). Cells treated with Ranpirnase recombinants exhibited a significant initial decrease in OD550 readout compared to PBS-treated cells, but reached control levels at 96 hours post-transfection, except for eGFP and p53 expression, which showed significant growth suppression. The expression of eGFP and RFP increased in cells treated with rBS-RNase, rRanpQ, and rRanp over time and the data obtained at 72 hours is presented in Supplementary Figure 4. These results suggest that co-expression of rBS-RNase, Bax, rRanpQ, and rRanp induced cytostatic effects during the initial three days.

Cytotoxicity of Ranpirnase recombinants was assessed using a Clonogenic assay. A549, NCI-H322, and NCI-H1299 cells were transfected with a mixture of pcDNA3.1 DNA and recombinant Ranpirnase expression plasmid DNAs at a 1:9 mass ratio. Transfected cells were cultured under G418 selection until colony formation. Representative results of three cell lines are shown in Supplementary Figure 5. Colony formation rates for the tested recombinant genes were analyzed (Fig 4d). Co-expression of hBax and p53 genes exhibited varying levels of inhibition to all cells, with discrepancy based on cell type. rBS-RNase, rRanpQ, and rRanp had no inhibitory effect on colony formation but increased the expression level of co-transfected eGFP and RFP genes (Supplementary Figure 4).

### 3.5. Toxicological Impact of rRanp Co-Expression

Hematological features and pro-inflammatory cytokines (TNF-α and IFN-γ) were analyzed in mouse blood after intramuscular administration of mѱCovS-LNP and mѱCovS-rRanp-LNP at a 15 µg mRNA dose. Mice did not exhibit significant weight loss (Supplementary Figure 6). Most blood parameters remained within normal ranges, except for decreased lymphocyte and platelet counts, elevated ALT levels, and transient increases in TNF-α and IFN-γ (Fig. 5ac-f). These observations suggest lymphocyte homing, mild liver injury, and inflammation induced by tested LNPs. Co-expression of rRanp showed no additional impact on ALT, TNF-α, and IFN-γ despite enhanced protein expression.

**Fig.5.**
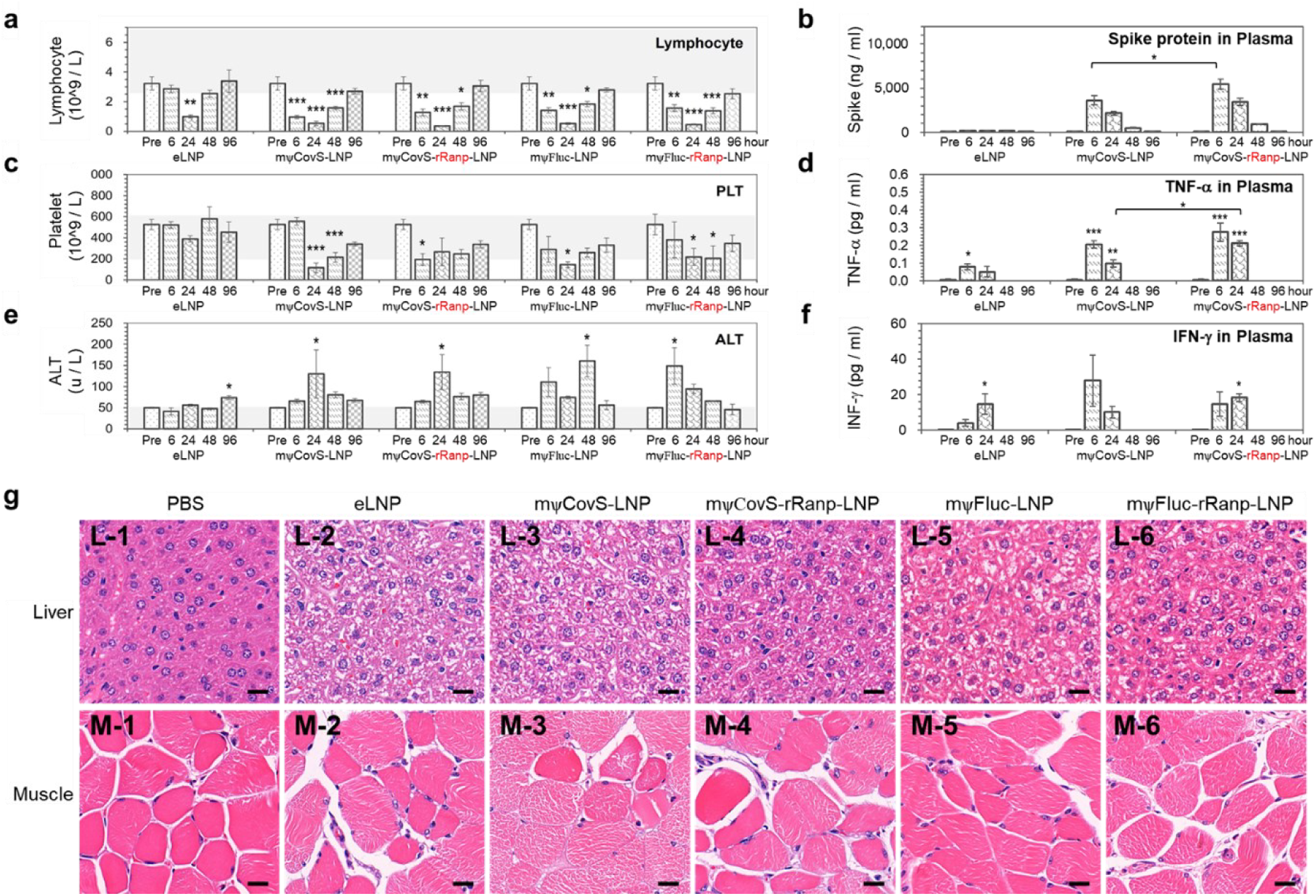
Alterations in Hepatotoxic and Immunological Biochemical Biomarkers in Balb/c Mice. Following Intramuscular Administration of CovS mRNA and CovS-rRanp mRNA encapsulated in LNP. LNP without mRNA (eLNP) and PBS were used as negative controls. (a) Lymphocyte count, (b) Spike protein levels, (c) Platelet count (PLT), (d) TNF-α levels, (e) Alanine aminotransferase (ALT) levels, and (f) IFN-γ levels in plasma samples were measured at pre-treatment and 6, 24, 48, and 96 hours post-administration. Physiological ranges of analytes in Balb/c mice are illustrated as gray boxes. (g) Representative H&E staining images from liver and muscle tissue. Bar size: 20 µm. Statistical significance between cohorts was determined using One-Way ANOVA with Bonferroni post-hoc tests (\**p* <0.05, \*\**p* < 0.01 and \*\*\**p* < 0.001). Error bars represent mean ± SEM.

Increased Spike protein levels were detected in the plasma of the mѱCovS-rRanp-LNP group compared to the mѱCovS-LNP control group by using a quantitative ELISA (Fig 5b). However, there was a discrepancy between protein levels and Fluc flux measured by BFI, which is likely attributed to the transmembrane nature of the Spike protein and divergent clearance mechanisms in blood and tissues (liver and muscle). Sustained production of Spike protein is suggested to result from rRanp-enhanced mRNA expression rather than solely relying on protein half-life prolongation.

To examine inflammation and toxicity, histological analysis was performed on mouse liver samples using H&E staining two days post-injection. Despite the increased production of Spike and Fluc proteins in the presence of rRanp, comparable cytoplasmic vesicular changes were observed in liver tissues for the eLNP group, the mRNA-LNP groups and rRanp-enhanced mRNA-LNP groups. Representative H&E images depicting these changes are presented (Fig 5g, liver). Muscle samples obtained from the injection sites were also evaluated and blindly graded by using a modified Allred scoring system based on the criteria outlined in Table 1. Representative H&E images are displayed (Fig 5g, muscle). A comparable level of moderate interstitial edema, local inflammation, and partial myofiber degeneration was observed for eLNP and all mRNA-LNP treatments. Necrosis of a few myofibers was found in samples of the mѱCovS-mRNA group regardless of rRanp status. The analysis suggests that host cells with rRanp co-expression increased tolerance to potentially deleterious proteins.

## 4. Discussion

Our study provides compelling evidence that the co-expression of rRanp significantly enhances transgene expression levels both in vivo and in vitro, while concurrently prolonging the expression duration. Notably, rRanp exhibits a gene expression enhancement effect on mѱ-modified mRNA, thereby improving the immunogenicity of mѱ-modified mRNA vaccines. Furthermore, rRanp has been observed to significantly enhance the expression levels of unmodified mRNA, offering an expression enhancement alternative when nucleoside modification is not feasible, such as with circular RNA. Importantly, co-expression of rRanp does not exacerbate adverse effects in the liver and muscle when compared to their mRNA counterparts. Additionally, histological assessment has revealed an increased tissue tolerance to potentially deleterious protein transcripts in the presence of rRanp. These findings highlight the potential value of rRanp as a novel expression enhancement tool for therapeutic applications, especially in mRNA-based therapies, while prioritizing safety and tolerability for recipients.

The expression-enhancing activity of rRanp demonstrates a unique mode of action that differentiates it from Ranpirnase. The removal of N-terminal modifications eliminates the ability of rRanp to penetrate tumor cells, which is a critical feature of Ranpirnase for exerting antitumoral cytotoxicity^26, 39, 40^. However, rRanp retains its ribonucleolytic activity^41^ and RI evasion ability, leading to a novel biological function characterized by lower antigenicity and cytotoxicity^26, 42, 43^.

Ranpirnase primarily functions through its tRNA-specific ribonucleolytic activity^19, 24^. The degradation of tRNA disrupts host metabolic activity, inducing a cytostatic state in cells^19^. Consequently, the degradation of both mRNA and protein transcripts is slowed down. Furthermore, Ranpirnase-induced tRNA degradation seems to coincide with an increase in tRNA turnover and the induction of new tRNA synthesis, resulting in a steady-state of tRNA levels within host cells^19^. Our metabolic activity analysis results support the observation that the co-expression of rRanp produces cytostatic effects rather than cytotoxic effects on host cells. This reduced biological activity is advantageous for maintaining protein equilibration when multiple genes are co-expressed. For instance, lower levels of metabolic activity may facilitate the efficient assembly of virus-like particles by equilibrating the quantities of multiple surface antigens and core proteins translated. The mRNA-1010 elicited higher HAI titers than a standard dose, inactivated seasonal influenza vaccine for influenza A strains, but comparable HAI titers for influenza B strains^44^, which could benefit from the rRanp enhancement potentially.

In addition to its effects on tRNAs, Ranpirnase also targets miRNA precursors, which bear structural similarities to tRNAs^20, 21^. Ranpirnase degradation of miRNA precursors leads to the downregulation of high-abundance miRNAs, particularly immune regulatory factors such as miR-155 and miR-21, thereby reducing innate immune responses^20^. This cleavage disrupts the maturation of miRNAs/siRNAs and diminishes the innate immune reactivity of cells. The miRNA-mediated RNA silencing mechanism is one of the key pathways for mRNA degradation, and Ranpirnase-mediated cleavage weakens this RNA decay process, ultimately increasing mRNA stability. Moreover, Ranpirnase plays a pivotal role in suppressing the innate immune response by preventing the translocation of NFκB to the nucleus^45, 46^.

With its compact size of 104 amino acids, rRanp gene can be seamlessly fused with target genes of interest or co-transfected alone, compatible with various delivery technologies. Furthermore, the implementation approaches for the rRanp gene are diverse, as it can be co-transfected with modified mRNA, unmodified mRNA, or plasmid DNA in existing protocols. This simplifies the experimental setup and eliminates the need for complex modifications or additional steps.

Although the safety of rRanp enhancement technology has been rigorously evaluated and found to have a favorable safety profile, further studies employing additional animal models, especially the non-human primates, and extended study durations are necessary before clinical translation can be pursued. An important challenge that remains to be addressed is the antigenicity of rRanp at repeated administration. Ranpirnase, which closely mimics the structure of RNase A and is highly conserved^47^, demonstrates very low antigenicity in mice^43^. In the absence of N-terminal modification, rRanp may possesses even lower antigenicity than Ranpirnase^48^. However, further investigations are required to confirm these findings. Additionally, co-transfection of rRanp resulted in elevated plasma levels of Spike protein, a transmembrane protein indeed. The detected increase in Spike protein in plasma did not align with the Fluc flux measurements obtained by using BFI. Future studies should consider employing a secretory peptide as an alternative detection method. Nevertheless, given the well-established safety profile of the mRNA platform, the application of rRanp enhancement technology in various fields, including veterinary use, appears imminent.

## Author Contributions

All authors confirmed they have contributed to the intellectual content of this paper with final approval of the publication. K.X., and H.Y conceived the concept and prepared the manuscript. X.W., C.H., Y.X.L., H.Y., and K.X. designed the experiments and analyzed data. K.X. performed pathological analysis and Allred scoring report. X.P., C.H., X.W., Y.X.L., and L.C design and carried out the animal experiments.

## Supporting information

Manuscript

## Acknowledgements

The authors thanked Dr Qinglong Hu, Tucson Pathology Associates, PC Carondelet Saint Joseph Hospital, Tucson, AZ 85711 USA., for his assistance in pathological analysis. Jian Yang and Lindan Diao for their technical assistances.

