## Supplementary material for "Recombinant Ranpirnase enhances the expression of co-transfected mRNA by modulating cellular metabolic activity": Manuscript

3: Animal experimental center of West China hospital, Sichuan University, Chengdu, 610041, China.

[a] These authors contributed equally to this work.

[b] Corresponding author:

 (K.Xu)

 (H.Yin)

This PDF file includes:

1. DNA dose determination for metabolic activity assay on NCI-H1299 cells

Supplementary Figure 1. The sequence comparison of Ribonucleases and their Recombinants3.

Supplementary Figure 2. Bioluminescent images for mice treated with rRanp-enhanced Fluc

Supplementary Figure 3. Influence of co-transfection of Fluc, rRanp, and p53 DNAs on the expression of pEGFP-c3 in A549, and NCI-H322 cells.

Supplementary Figure 4. Influence of Ribonuclease Recombinants on Metabolic Activity

Supplementary Figure 5. Clonogenic assay on cultured cells.

Supplementary Figure 6. Weight changes of mice after treated with 15 μg of mRNA-LNP.

### 1. DNA dose determination for metabolic activity assay on NCI-H1299 cells

To determine the appropriate DNA dose with minimal cytotoxic effects, a pilot study was conducted.

The metabolic activity of NCI-H1299 cells was evaluated using an XTT assay after treatment with recombinant ribonucleases. The cells were seeded in 96-well microplates at a density of 5,000 cells per well and initially transfected with DNA:DL complexes containing 425, 212, and 42.5 ng of pEGFP-c3 DNA, separately. The metabolic activity of the treated cells was compared to that of PBS-treated cells for normalization purposes. After 96 hours post-transfection, the cell survival rates were determined to be 16.01%, 28.54%, and 94.24% for the pEGFP-treated cells, respectively, indicating the baseline level of cytotoxicity associated with the transfection complexes. Based on the results of the pilot study, a DNA dose of 42.5 ng per well was chosen for subsequent evaluations.

**Supplementary Figure 1.** The sequence comparison of Ribonucleases and their Recombinants

```
    Name                                   Amino Acid sequence
                                                    1
Ranpirnase Precursor:      mfpkfsfllifavvlslthkslcqdWLTFQKKHITNTRDVDCDNIMSTNLFHCKDKNTFIYSR  63
Ranpirnase (mature):                          +pyrQDWLTFQKKHITNTRDVDCDNIMSTNLFHCKDKNTFIYSR  39
                                                 -1 1
rRanp:                                           MSDWLTFQKKHITNTRDVDCDNIMSTNLFHCKDKNTFIYSR  40
rRpK31R:                                         MSDWLTFQKRHITNTRDVDCDNIMSTNLFHCKDKNTFIYSR  40
rRanpQ:                                          MQDWLTFQKKHITNTRDVDCDNIMSTNLFHCKDKNTFIYSR  40


Ranpirnase Precursor:     PEPVKAICKGIIASKNVLTTSEFYLSDCNVTSRPCKYKLKKSTNKFCVTCENQAPVHFVGVGSC  126
Ranpirnase (mature):      PEPVKAICKGIIASKNVLTTSEFYLSDCNVTSRPCKYKLKKSTNKFCVTCENQAPVHFVGVGSC  104

rRanp:                    PEPVKAICKGIIASKNVLTTSEFYLSDCNVTSRPCKYKLKKSTNKFCVTCENQAPVHFVGVGSC  105
rRpK31R:                  PEPVKAICKGIIASKNVLTTSEFYLSDCNVTSRPCKYKLKKSTNKFCVTCENQAPVHFVGVGSC  105
rRanpQ:                   PEPVKAICKGIIASKNVLTTSEFYLSDCNVTSRPCKYKLKKSTNKFCVTCENQAPVHFVGVGSC  105


Name                                       Amino Acid sequence
                           1
Amphinase-1 (mature):      REWEKFKTKHITSQSVADFNCNRTMNDPAYTPDGQCKPVNTFIHSTTGPVKEICRRATGRVNK  63
                           SSTQQFTLTTCKNPIRCKYSQSNTTNFICITCRDNYPVHFVKTGKC                  115
                        -1 1
rAmph-1:                  MREWEKFKTKHITSQSVADFNCNRTMNDPAYTPDGQCKPVNTFIHSTTGPVKEICRRATGRVNK  64
                           SSTQQFTLTTCKNPIRCKYSQSNTTNFICITCRDNYPVHFVKTGKC                  116

                           1
BS-RNase (mature):         KESAAAKFERQHMDSGNSPSSSSNYCNLMMCCRKMTQGKCKPVNTFVHESLADVKAVCSQKKV  63
                           TCKNGQTNCYQSKSTMRITDCRETGSSKYPNCAYKTTQVEKHIIVACGGKPSVPVHFDASE   124
                        -1 1
rBS-RNase:                MKESAAAKFERQHMDSGNSPSSSSNYCNLMMCCRKMTQGKCKPVNTFVHESLADVKAVCSQKKV  64
                           TCKNGQTNCYQSKSTMRITDCRETGSSKYPNCAYKTTQVEKHIIVACGGKPSVPVHFDASE   125

                           1
RNase I (mature):          KESRAKKFQRQHMDSDSSPSSSSTYCNQMMRRRNMTQGRCKPVNTFVHEPLVDVQNVCFQEKVT 63
                           CKNGQGNCYKSNSSMHITDCRLTNGSRYPNCAYRTSPKERHIIVACEGSPYVPVHFDASVEDST 128
                        -1 1
rRNase I:                 MKESRAKKFQRQHMDSDSSPSSSSTYCNQMMRRRNMTQGRCKPVNTFVHEPLVDVQNVCFQEKVT 64
                           CKNGQGNCYKSNSSMHITDCRLTNGSRYPNCAYRTSPKERHIIVACEGSPYVPVHFDASVEDST 129
```

**Supplementary Figure 2.** Bioluminescent images for mice treated with rRanp-enhanced Fluc in different LNPs. Bioluminescent images of mice obtained using IVIS imaging at various time points (0.25, 1 to 15 days) following intramuscular injection of 5 μg of mψ-mRNA encapsulated in LNP and LNP$^{\oplus}$74 (n = 3 mice per group), either with or without rRanp.

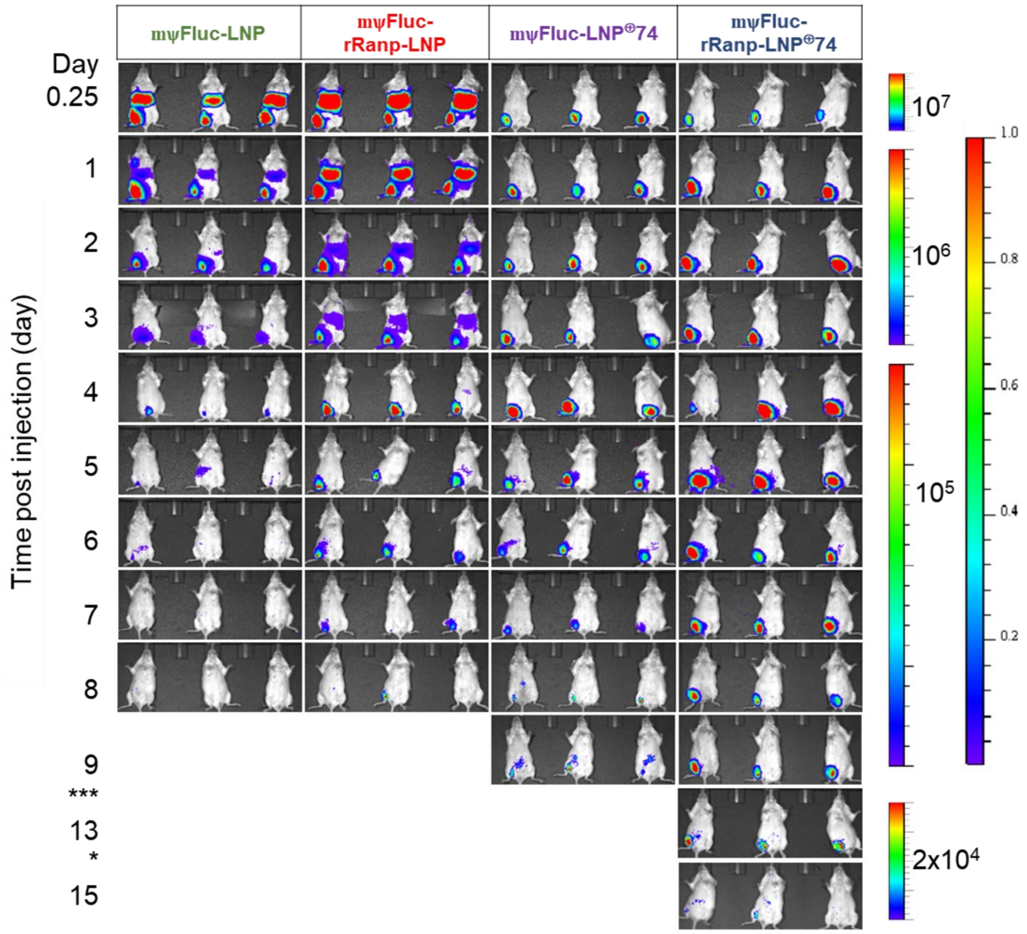


**Supplementary Figure 3.** Influence of co-transfection of Fluc, rRanp, and p53 DNAs on the expression of pEGFP-c3 in A549, and NCI-H322 cells.

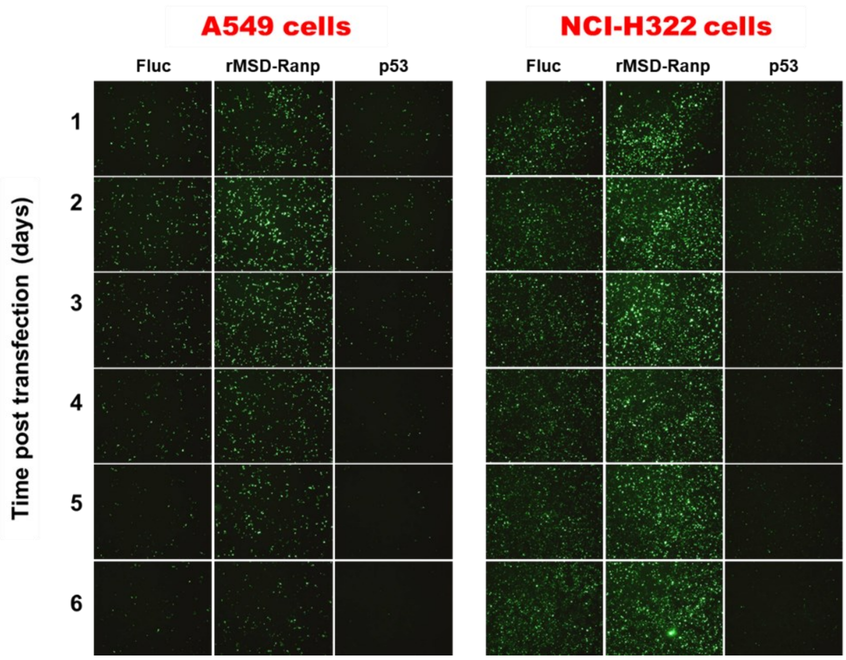

**Supplementary Figure 4.** Influence of Ribonuclease Recombinants on Metabolic Activity of NCI-H1299 cells 72 hours after treatment. Cell viability was measured with an XTT assay.

| Plasmid | CMV-eGFP | CMV-BS-RNase | CMV-hBax | CMV-p53 | CMV-rRanpQ | CMV-rRanp |
|---|---|---|---|---|---|---|
| CMV-RFP | 4.25 ng | 4.25 ng | 4.25 ng | 4.25 ng | 4.25 ng | 4.25 ng |
| CMV-eGFP | 38.25 ng | 4.25 ng | 4.25 ng | 4.25 ng | 4.25 ng | 4.25 ng |
| CMV-BS-RNase | | 34 ng | | | | |
| CMV-hBax | | | 34 ng | | | |
| CMV-p53 | | | | 34 ng | | |
| CMV-rRanpQ | | | | | 34 ng | |
| CMV-rRanp | | | | | | 34 ng |
| Cell Viability | 70.56% | 74.28% | 86.15% | 62.98% | 81.45% | 71.05% |

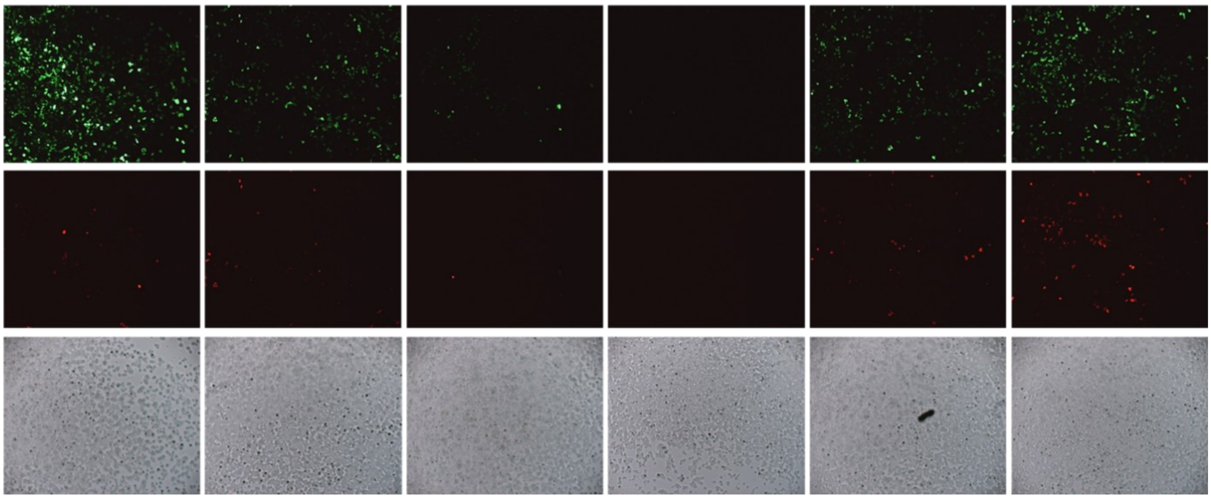


**Supplementary Figure 5.** Clonogenic assay on cultured cells.

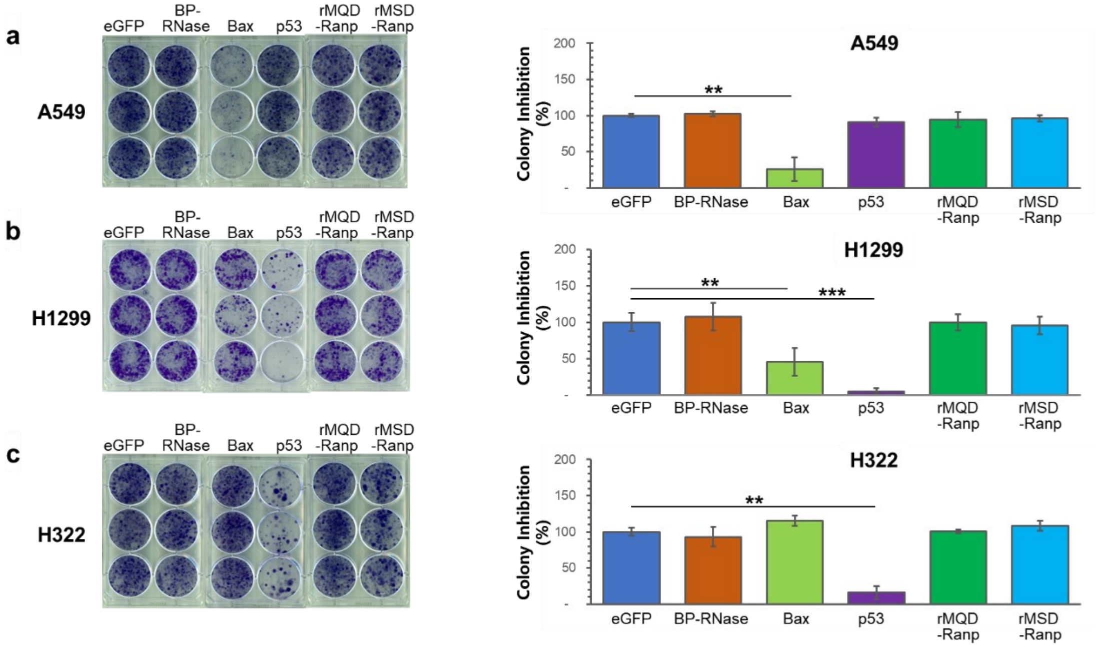


**Supplementary Figure 6.** Weight changes of mice after treated with 15 μg of mRNA-LNP.

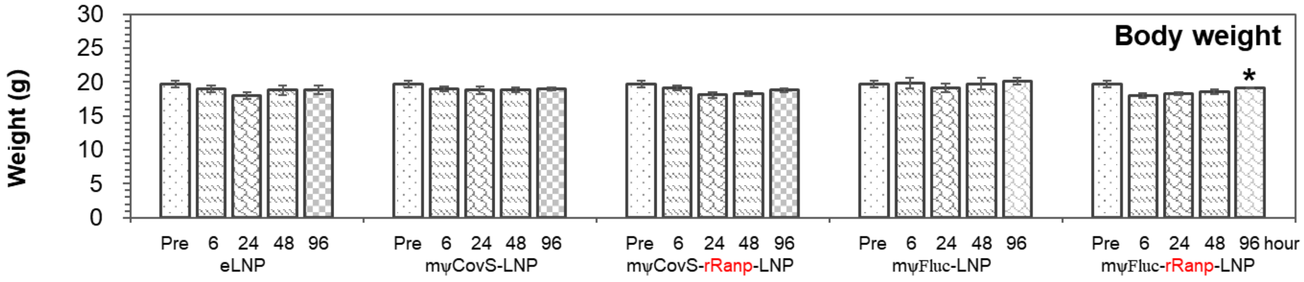

Body weight
Weight (g)
30
25
20
15
10
5
0
*
Pre 6 24 48 96
eLNP
Pre 6 24 48 96
mψCovS-LNP
Pre 6 24 48 96
mψCovS-rRanp-LNP
Pre 6 24 48 96
mψFluc-LNP
Pre 6 24 48 96 hour
mψFluc-rRanp-LNP
